# Adhirons are efficient tools to guide antiviral ligand discovery

**DOI:** 10.1101/2025.05.13.653658

**Authors:** Alex J. Flynn, Alex Derry, Oliver Debski-Antoniak, Stephen P. Muench, Juan Fontana, Katie Simmons

## Abstract

Adhirons are small, antibody-like proteins expressed in bacteria that bind to biological targets with high affinity. Since their discovery, Adhirons have been isolated against over 350 targets, and they can be used for many research applications. However, their therapeutic application is at present in its early stages, hence the interest in developing novel small molecules to mimic their effect. We developed a pipeline for small molecule discovery where Adhirons act as starting templates in a ligand-based virtual screening workflow. For proof of concept, we used Adhirons against two viral systems: Crimean-Congo Haemorrhagic Fever Virus and Influenza A Virus. In both cases, a hit was identified from the mimics using biophysical and cell-based assays, highlighting how Adhirons, commercial libraries and structural approaches can be combined to accelerate small molecule inhibitor discovery of viruses.

## Introduction

As evidenced by the recent SARS-CoV-2 pandemic, viral diseases can be lethal, causing lasting ill health and world-wide disruption. While vaccination is an effective way to prevent viral infections, antivirals are also important tools for treating severe cases. However, developing an antiviral or any new drug is a long process [1]. To accelerate the drug development process, different methods for guiding small molecule discovery have been proposed.

Antibodies can be highly effective at targeting specific proteins and neutralising viral infection, but their use can be limited by poor pharmacokinetics and poor tissue penetration. One promising approach is for the development of therapeutic small molecules involves mimicking neutralising antibodies. Several examples of antibody-guided small molecule development exist in the literature. For example, a monoclonal antibody (mAb) binding region was exploited to develop a cyclic organic small molecule via a peptide mimetic intermediate molecule [2]. The resulting small molecule acted through the same pathways as the original template mAb. More recently, a broadly neutralising antibody targeting the hemagglutinin (HA) spike protein in influenza A virus (IAV) was used to generate a small molecule using high throughput screening [3]. The resulting small molecule was similarly broadly acting, bound to the same location on HA as the template antibody, and had the benefit of being orally bioavailable. With more available high-resolution protein structures than ever, improved ligand-based virtual screening algorithms and larger commercially accessible small molecule libraries, antibody-guided small molecule development could extend beyond just the few published examples. However, the large binding interface of antibodies mean other biopharmaceuticals may be better suited as templates for small molecule development.

Adhirons are antibody-like proteins based on a constant 91 amino acid scaffold protein that constrains two randomised nine amino acid variable regions for molecular recognition. Adhirons are extremely stable at high temperatures and easy to express and purify from *E. coli*, allowing for large scale production at low costs. A phage display library containing an even distribution of each of the 19 amino acids (excluding cysteines) in the variable regions, provides >3×10^10^ unique Adhirons that can be screened against a molecule of interest, allowing the selection of specific Adhirons with high affinity against any given target. Typically, Adhiron selection consists of three panning rounds of phage display and phage ELISA against a target protein. This approach has allowed the isolation of Adhirons against more than 350 targets. At ∼12 kDa, Adhirons are not only much smaller than antibodies, but they can also access smaller pockets on the target surface due to their smaller binding regions. Recently, the loop residues in an Adhiron responsible for binding and inhibiting KRAS showed high structural similarity to a small molecule inhibitor that bound at the same site, leading to speculation that the Adhiron could be used to develop small molecule mimics in the future [4]. Furthermore, Adhirons binding to Crimean-Congo haemorrhagic fever virus (CCHFV) [5] and IAV [6] have been identified, making them potential templates to develop a method of Adhiron-guided small molecule discovery.

CCHFV is the causative agent of Crimean-Congo Hemmorhagic Fever (CCHF) disease. Symptoms of CCHF include fever, nausea, organ failure and haemorrhage and CCHF has a fatality rate estimated at 5-30% [7] . CCHFV is spread by *Hyalomma* ticks, which bite humans and release the virus into the bloodstream. The vast tick distribution across Africa, Asia, the Middle East and Southern and Eastern Europe leads to regular CCHFV outbreaks in these regions, making CCHFV the most wide-spread tick-borne virus and the only biosafety level 4 (BSL4) virus that is endemic in Europe [8]. Its severity and wide distribution caused the world health organisation to rank CCHF in the top priority diseases for research and development [9].

Treatment for CCHFV infection is currently limited to Ribavirin, a broadly-acting guanosine analogue which has several proposed modes of action, including direct inhibition of the viral RNA dependant RNA polymerase (L), inhibition of enzymes involved in guanosine metabolism and incorporation into viral RNA products by L polymerase, which introduces lethal mutations that result in defective virus particles [10]. However, although *in-vitro* assays and clinical observational studies indicate ribavirin treatment inhibits CCHFV infection, a systematic review found insufficient evidence to demonstrate the efficacy of Ribavirin in treating CCHF [11]. Favipiravir is another guanosine analogue that has similar modes of action to ribavirin and causes lethal mutations in CCHFV, but it has not yet been approved for human use [12]. Furthermore, mAbs against Gc and Gn protected mice from a lethal CCHFV challenge [13]. Three of these mAbs were broadly neutralising against glycoproteins from a range of CCHFV strains, preventing CCHFV from entering the required cells, but these mAbs are yet to be tested in humans.

The need for new reagents to treat and diagnose CCHFV was the basis for the development of an Adhiron, named NP-Adhiron, which was directed against the CCHFV nucleoprotein (NP), a viral protein involved in packing the viral RNA that is essential at many stages of the viral replication cycle [5]. Importantly, NP-Adhiron inhibited CCHFV replication when tested using a CCHFV mini-genome system, but its inability to enter cells hindered its use as a therapeutic. However, given NP-Adhiron binds to CCHFV NP with high affinity and strongly inhibits CCHFV replication, we reasoned that the region of NP recognised by NP-Adhiron was a good target for small molecule inhibitors, and that NP-Adhiron loops were good templates to use for identifying such small molecules.

A second Adhiron, targeting the HA spike protein from IAV, was isolated as a further template for small molecule identification. IAV causes ‘the flu’, a respiratory illness with symptoms including a sudden high temperature, coughing, headache and fatigue. An estimated 300,000-500,000 deaths are attributed annually to seasonal influenza epidemics [14]. The constant emergence of new strains necessitates annual vaccination programs and leads to the resistance of strains to antiviral therapeutics. Of particular concern is the highly pathogenic avian influenza (HPAI) H5N1, which has shown an expanding host range, infecting not only poultry but also wild and domestic mammals [15]. Its continued circulation increases the risk of spillover into humans, raising fears of a potential pandemic with high mortality rates, especially given its historical case fatality rate exceeding 50% in humans and recent detections in mammalian hosts.

There are several small molecules approved for the treatment of IAV, but all have problems related to viral resistance. The first anti-influenza drugs to be approved were the M2 channel inhibitors amantadine and rimantadine, but widespread resistance to them means they are no longer in use [16]. Currently, the first-line treatments are NA glycoprotein inhibitors such as zanamivir, peramivir and oseltamivir, which are effective at reducing illness severity and duration. However, some circulating strains are starting to show resistance to these compounds [17]. Additionally, baloxavir is a polymerase inhibitor that has been approved for use in several countries. Unfortunately, a study found that 10% of patients with influenza that were treated with baloxavir developed a mutation associated with resistance [18]. As of yet, there are no approved inhibitors of the HA glycoprotein, but various compounds targeting HA have demonstrated inhibition of IAV infection in mice [3], indicating that HA is a promising target that could be investigated further.

Here we demonstrate how the structural features of anti-viral Adhirons can be used to develop small molecule mimics of their binding regions. We demonstrate that molecules identified in this way can bind to the viral target protein and inhibit activity with micromolar potency. Subsequently, we propose a workflow for Adhiron-based small molecule discovery and characterisation which could be applied to a range of viral targets and other disease areas.

## Results

### Small molecule mimics can be identified using structural information of an Adhiron bound to its target

A typical Adhiron loop contains nine amino acids, weighing approximately 990 Daltons-which is much larger than most drug-like molecules found in commercially available libraries. To define the region of the Adhiron loops binding to CCHFV, the X-ray structure of the complex CCHFV NP-Adhiron:NP [5] was analysed for the key interactions to retain with our virtual screening efforts. The NP-Adhiron binds within a shallow groove on the head domain of CCHFV NP (Fig. 1A), which was proposed to be the site of NP oligomerisation [5]. Visual inspection of the complex suggests that NP-Adhiron loop two is positioned near to the surface of NP, whereas loop one does not interact with NP (Fig. 1B). In particular, the residues in the second half of loop two, positions 106-110, are in closest proximity to NP, with F107 and W108 residing in a groove between two helices (Fig. 1B). To quantify the binding effect of each residue within the binding pocket, we employed PDBePISA [19], to measure the buried surface area/accessible surface area (BSA/ASA), which is the percentage of the surface area that is buried upon binding; and the solvation energy, a measure of the contribution that polar and hydrophobic interactions make towards binding. A positive solvation energy indicates interactions are based on the hydrophobic effect and a negative value indicates the interactions are due to the polar interactions between residues. Calculating the BSA/ASA identified that four residues (Y103, F107, W108 and D110) in loop two are buried >70% upon binding, while only R73 from loop one is similarly buried (Fig. 1C). Therefore, NP-Adhiron loop two is the primary determinant of binding, with residues 106-110 forming its core interaction site with CCHFV NP. Solvation energy analysis highlights that the chemical basis for interactions varies throughout loop two. For example, binding at Y103, F107 and W108 is driven by the hydrophobic effect, but binding at D106 and D110 is driven by polarity (Fig. 1C). Hydrogen bonding interactions with CCHFV NP are formed by Y103, D106, F107 and W108 and a salt bridge involving D110 in NP-Adhiron (Fig. 1C). Overall, this analysis allowed us to identify the second half of loop two from NP-Adhiron (D106-D110) as the core site of binding, and that a combination of interactions driven by polarity and hydrophobicity are important for CCHFV NP:NP-Adhiron binding. Therefore, NP-Adhiron loop two could serve as a valuable template for small-molecule design.

**Figure 1:**
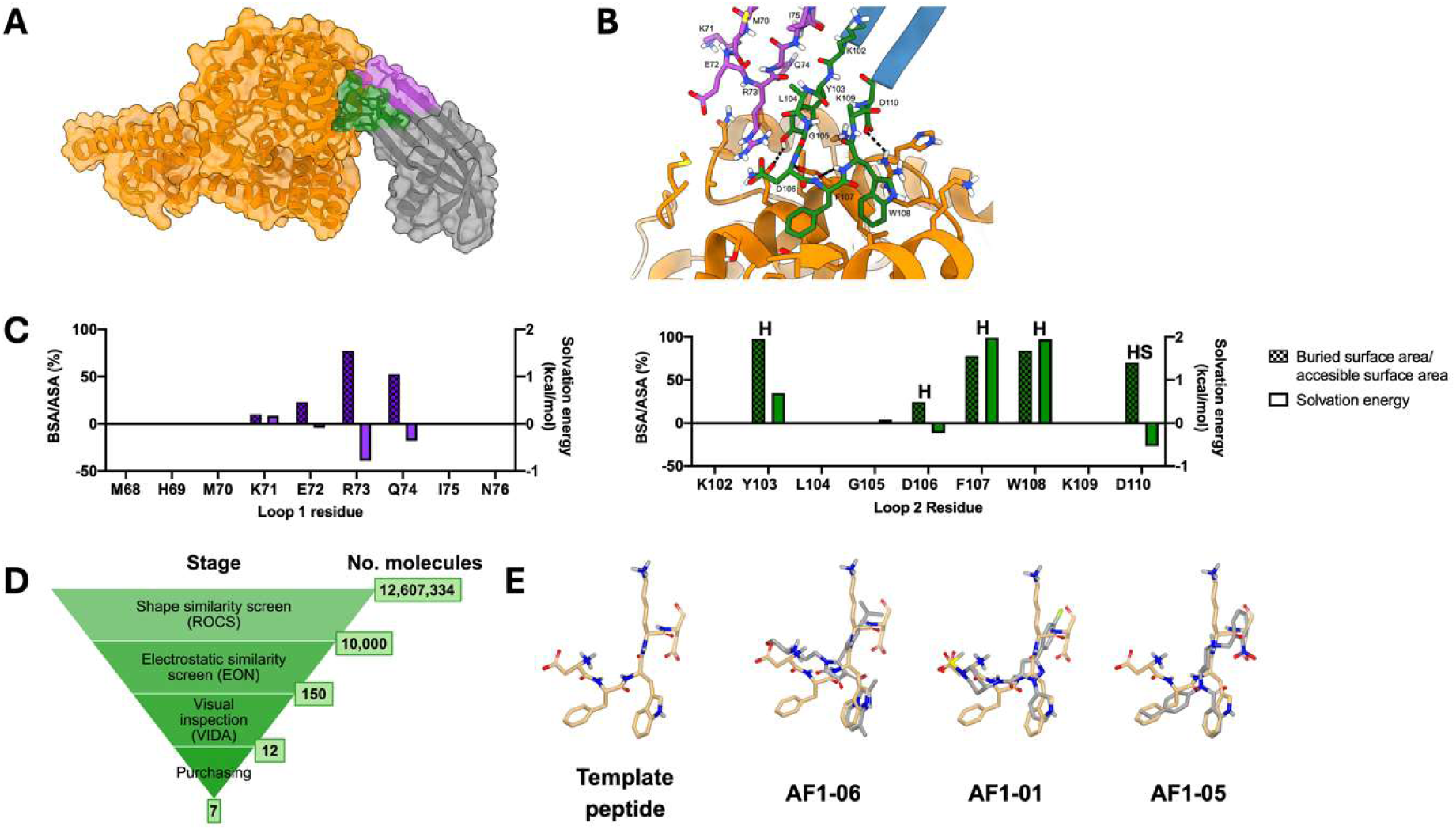
A ligand-based virtual screening workflow identified small molecules that mimic the binding region of an Adhiron (NP-Adhiron) that inhibits Crimean Congo haemorrhagic fever virus (CCHFV). A) The overall 3D structure of CCHFV NP bound to NP-Adhiron. Loop 1 is purple and loop 2 is green. B) The binding site of CCHFV NP to NP-Adhiron with Adhiron loop residues labelled. C) PDBePISA analysis of NP-Adhiron:NP crystal structure showing buried surface area/accessible surface area and solvation energy for each NP-Adhiron residue. Residues where hydrogen bonds or salt bridges were identified are labelled with H and HS respectively. D) Virtual screening workflow used to identify small molecule mimics of the Adhiron loop from the eMolecules database of 12.6 million compounds. E) Example compounds from the virtual screen. The NP-Adhiron DFWKD template is shown as beige sticks and the compounds are shown as grey sticks.

The D106-D110 (DFWKD) pentapeptide from loop two was computationally excised from the 3D structure and used as the template in a ligand-based virtual screening workflow. First, compounds in the eMolecules database were screened for those with high shape similarity for the pentapeptide using ROCS (ROCS 3.6.2.3. OpenEye, Cadence Molecular Sciences, Santa Fe, NM) [20,21], and the top matches were screened for electrostatic similarity to the template using EON (ROCS 3.6.2.3. OpenEye, Cadence Molecular Sciences, Santa Fe, NM) (Fig. 1D). The top scoring compounds from this process underwent visual inspection and promising compounds were identified such that they mimicked some of the pentapeptide main chain and at least one side chain, and they were in the same orientation as pentapeptide. For future synthetic tractability and selectivity, compounds with some hydrophobic fused ring structures or flexible chains were avoided. Additionally, based on the CCHFV NP:NP-Adhiron interactions, we selected small molecules that mimicked at least one of the three regions involved in hydrogen bonding or the salt bridge involved in the interaction. Furthermore, compounds were selected to mimic some of the hydrophobic interactions with residues including F107 and W108. Overall, this resulted in 12 selected compounds of which seven were available from commercial sources. These were purchased and termed AF1-01 to AF1-07 (Fig. 1D and Supp. Fig 1).

Due to the size of the small molecules versus the pentapeptide, it was not possible to mimic the entirety of the template pentapeptide. For example, whereas compound AF1-06 mimicked the W108 side chain through an imidazopyridine ring, it lacked the functional groups to mimic the acidic side chains of D106 and D110 (Fig. 1E). The acidic side chain of D106 of compound AF1-01 mimicked the sulfone group but the pyridine ring was a poor mimic of the W108 indole. Finally, compound AF1-05 mimicked the acidic side chain of D110 with a nitro group, but mimicked W108 poorly with a benzene ring. There were no compounds that mimicked K109 well, but this was deemed acceptable as the side chain of this residue is not directly involved in Adhiron-binding. (Fig. 1B and C). Overall, this analysis suggested that we would likely see reduced binding affinities for the small molecules compared to the template Adhiron, since the compounds only mimicked some but not all of the key binding interactions.

### Small molecule mimics of the CCHFV NP Adhiron demonstrate inhibition of viral replication in a mini-genome assay

Once a set of potential small molecule mimics of CCHFV NP-Adhiron were identified, we sought to characterise them *in vitro*. Initially, compounds were screened for cytotoxicity in the BSR-T7 cell line used for CCHFV mini-genome system [5]. Cells were incubated with 10 µM of the compounds and the resulting cell viability was assayed by measuring the intracellular ATP concentration, which rapidly declines in dying cells (Fig. 2A). Incubation with AF1-03 - AF1-07 resulted in a cell viability close to 100% and were therefore categorised as non-cytotoxic. Incubation with compounds AF1-01 and AF1-02 led to a cell viability of ∼70%, which was low but considered suitable for further evaluation.

**Figure 2:**
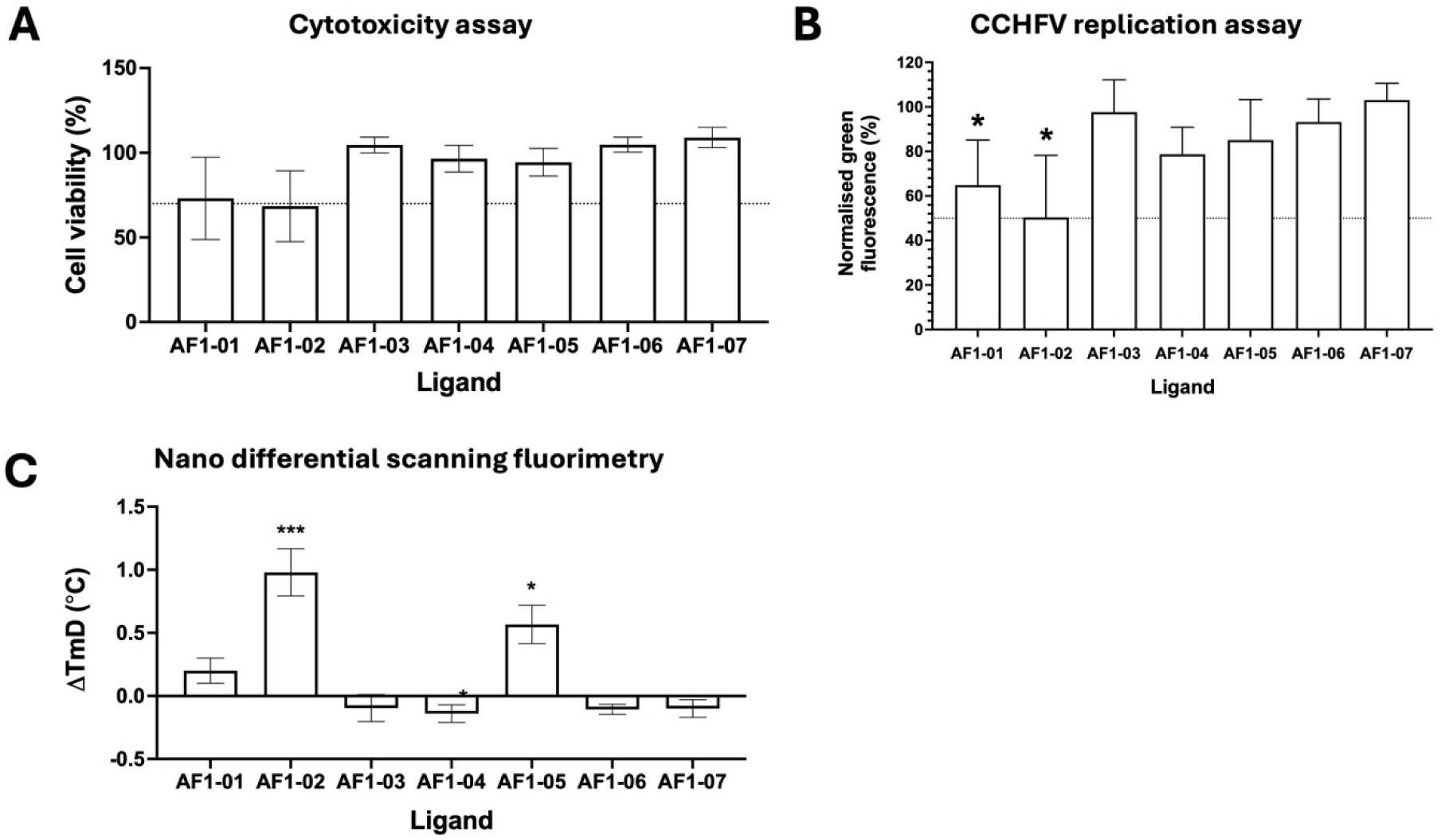
Screening of small molecule mimics of an Adhiron that binds CCHFV NP. A) Cytotoxicity assay in BSR-T7 cells with 10 μM compound (N=3). Results are normalised to a DMSO control. B) AF1 compound series screen at 10 μM against the CCHFV mini-genome system (MGS), a proxy for virus replication (N=3). Results normalised to the DMSO control. C) Screening the AF1 compound series at 200 μM using thermal shift assays by nanoDSF (N=3). NP + compound normalised to NP + DMSO control. Error bars indicate mean ± standard deviation. Tm shifts marked with their statistical significance according to a paired t-test. Significance levels for panels 2B and 2C: ns = no significance, * = p<0.05, ** = p<0.01, *** = p<0.001.

To test the inhibitory properties of these molecules, a mini-genome system was employed, which allows monitoring of viral replication via expression of a eGFP reporter gene [5]. Therefore, in the presence of replication inhibitors eGFP expression will be reduced. We optimised the method for transfecting the CCHFV mini-genome system from the one previously used to characterise the CCHFV NP-Adhiron [5]. Previously, plasmids for the mini-genome system were co-transfected with a plasmid for NP-Adhiron. However, this method was considered inappropriate for screening the small molecules as we reasoned that addition of compound could negatively affect the DNA transfection and produce a false positive result. Instead, cells were first transfected with the mini-genome plasmids, and after 24 hrs transfected cells were trypsinised and reseeded in the presence of the compounds. Using this approach, AF1-02 reduced GFP fluorescence to ∼50% (Fig. 2B) when tested at a concentration of 10 µM. Of note, the reduction in GFP expression was greater than the cell viability shown in the cytotoxicity experiment, suggesting that this compound inhibits CCHFV replication.

### Small molecule mimics of CCHFV NP-Adhiron bind to NP

Next, we aimed to test if the NP-Adhiron mimics bind to CCHFV NP using nano differential scanning fluorimetry (nanoDSF). Rather than measuring protein unfolding via an extrinsic fluorescence source (as occurs in standard DSF), nanoDSF measures changes in intrinsic protein fluorescence, based on the fact that the wavelength of fluorescence emitted by tryptophan residues shifts from 330 nm to 350 nm when a protein unfolds [22]. Measuring a positive shift in protein melting temperature (T_m_) in the presence of a ligand indicates the ligand has a stabilising effect on the protein via binding.

When the selected compounds were screened for binding to NP by nanoDSF, fluorescence at 330 nm and 350 nm was measured and the derivative of the ratio (F330/F350) was plotted against the temperature, generating melting peaks. Finally, the inflection point of each curve was measured as the T_m_ and the shift for each compound was calculated relative to the DMSO control (Fig. 2C). Both AF1-02 and AF1-05 generated a positive thermal shift for CCHFV NP of 0.98 °C and 0.57 °C respectively, indicating that they bind to CCHFV NP. Combined with the results from the mini-genome system assay, it suggests that AF1-05 binds CCHFV NP but is not biologically active, whereas AF1-02 both binds and inhibits CCHFV NP. Although AF1-02 possesses weak activity, this could be improved by rounds of structural optimization. These results demonstrate that Adhirons can be used as scaffolds to direct the identification of small molecule modulators.

### Adhiron small molecule mimics can be identified without structural information of Adhiron-target binding

While AF1-02 was identified as a mimic of a neutralising Adhiron for which there was a structure available, we also wanted to test if Adhiron mimics could be identified in the absence of a high-resolution Adhiron-target structure. Recently we described two Adhirons targeting HA from the pandemic strain A/Aichi/68 (H3N2) IAV, named Adhirons A5 and A31 [6]. They inhibit IAV infection via binding at the receptor-binding domain. However, the low resolution of the Adhiron-HA interface in the A5- and A31-bound HA cryo-electron microscopy maps prevented accurate determination of the exact Adhiron loop regions. Therefore, to generate mimics of A5 and A31, the Adhiron structures were first generated via homology modelling [23] using an anti-Bcl-XL Adhiron as a template [24]. In essence, this approach allowed us to model the Adhiron loop structures while restraining the Adhiron scaffold structure. Since no information was available on the specific residues responsible for binding, we computationally excised the entirety of both Adhiron loops from the homology models and used them as templates for the same ligand-based virtual screening workflow described for NP-Adhiron (Figure 3A).

**Figure 3:**
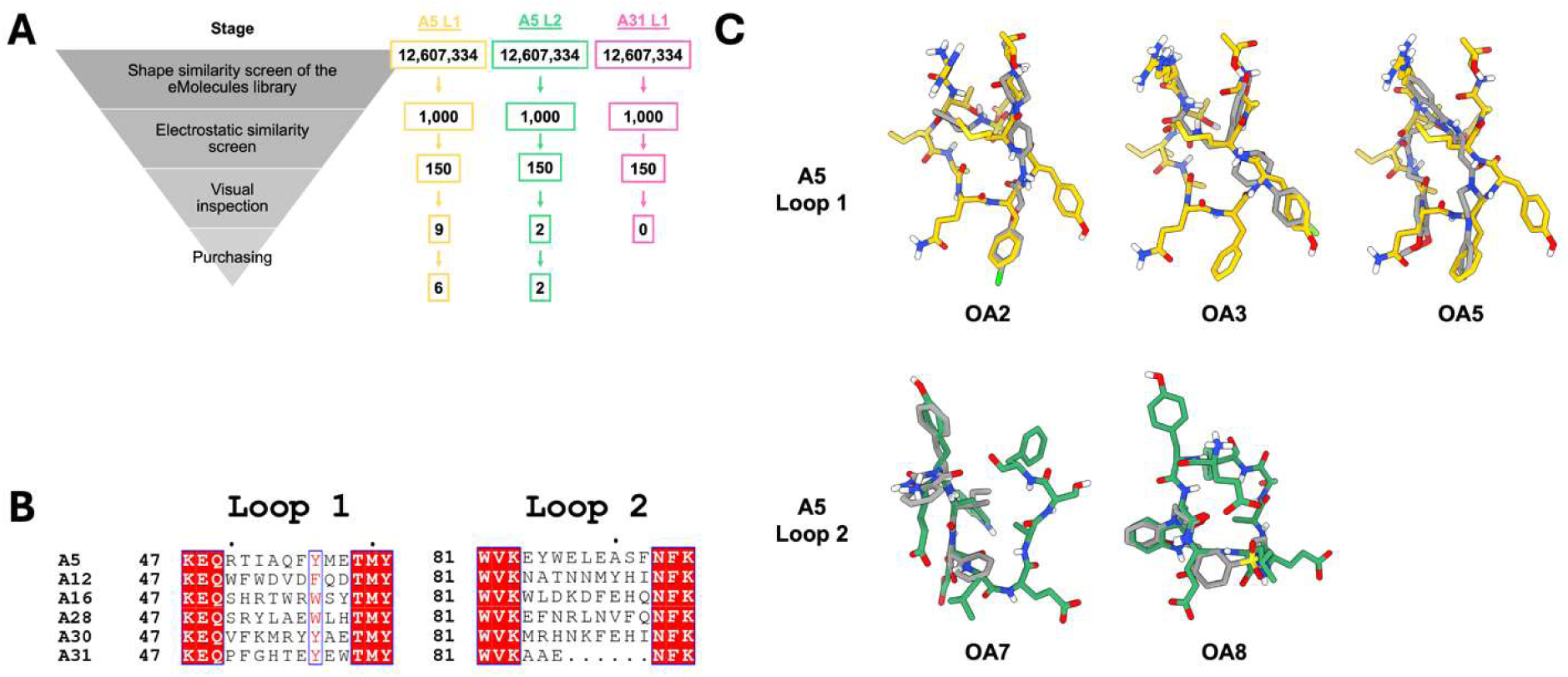
Development of small molecule mimics of neutralising Adhirons that bind inffuenza A virus hemagglutinin (HA). A) Ligand-based virtual screening workffow used to identify compounds from the eMolecules library that are structurally similar to either A5 loop 1 or loop 2. B) Sequence alignment of the two hypervariable loops in six Adhirons which bound HA. A5 and A31 were used as templates for their high potency of inhibition. Sequence identity indicated by a filled red box and sequence similarity indicated by red text. Image generated using the eSPRIPT server [25,2C]. C) Example small molecule mimics identified using the workffow in panel A. A5 loop 1, A5 loop 2 and small molecules shown as yellow, green, and grey sticks, respectively.

Due to a lack of structural information on the Adhiron residues responsible for binding HA, sequence alignment of six Adhirons which bind HA, including A5 and A31, was performed to identify amino acids likely involved in binding (Figure 3B) [6]. The alignment revealed that position seven of loop one was conserved in all Adhiron clones as an aromatic residue and was therefore mimicked in all identified molecules. We applied the same general rules as above: compounds should have the same shape and orientation as the loop, mimic at least one side chain and potential hydrogen bond donor or acceptor, and not contain large hydrophobic regions or long flexible chains. Eight compounds that mimic Adhiron A5 were selected-with compounds OA1-OA6 mimics of loop one and OA7 and OA8 mimics of loop two (Supp. Fig. 2). The large residues in A31 meant molecules did not accurately mimic the loop region, so no compounds were tested as mimics of this Adhiron.

As with the mimics of NP-Adhiron, the mimics of A5 loop one and two mimicked some functional groups well but not all. For example, OA2 contained a sulfonyl group that aligned with the carboxyl group of E58 and a fluorinated aromatic ring that overlaid well with F55. OA3 also had a fluorinated aromatic ring which mimicked the conserved Y56 well. Finally, OA5 had a tricyclic ring system that overlaid well with F55. Since no structural information was available on the binding of A5 to HA, compounds were purchased so that together they gave maximum coverage of each loop. This approach aimed to ensure that at least one compound would cover the region responsible for binding. For example, whereas OA2 mimics F55, OA3 mimics Y56 instead; and whereas OA7 covers the first third of loop two, OA8 covers the second third.

### Small molecule mimics of an Adhiron against IAV bind to HA and inhibit IAV infection in tissue culture

Prior to screening compounds for IAV inhibition in a TCID_50_ assay using Madin-Darby canine kidney (MDCK) cells, compounds were screened for cytotoxicity. Given several AF1 compounds were found to be cytotoxic at 10 µM, a range of concentrations for the OA series were screened (Fig. 4A). OA6 demonstrated the greatest cytotoxicity with a median lethal dose (LD_50_) of 3.6 µM and OA7 demonstrated the least cytotoxicity with a LD_50_ 390.9 µM. To avoid problems with compound toxicity, the highest concentration for each compound that led to >95% cell viability was selected as the maximum concentration for TCID_50_ assay screening.

**Figure 4:**
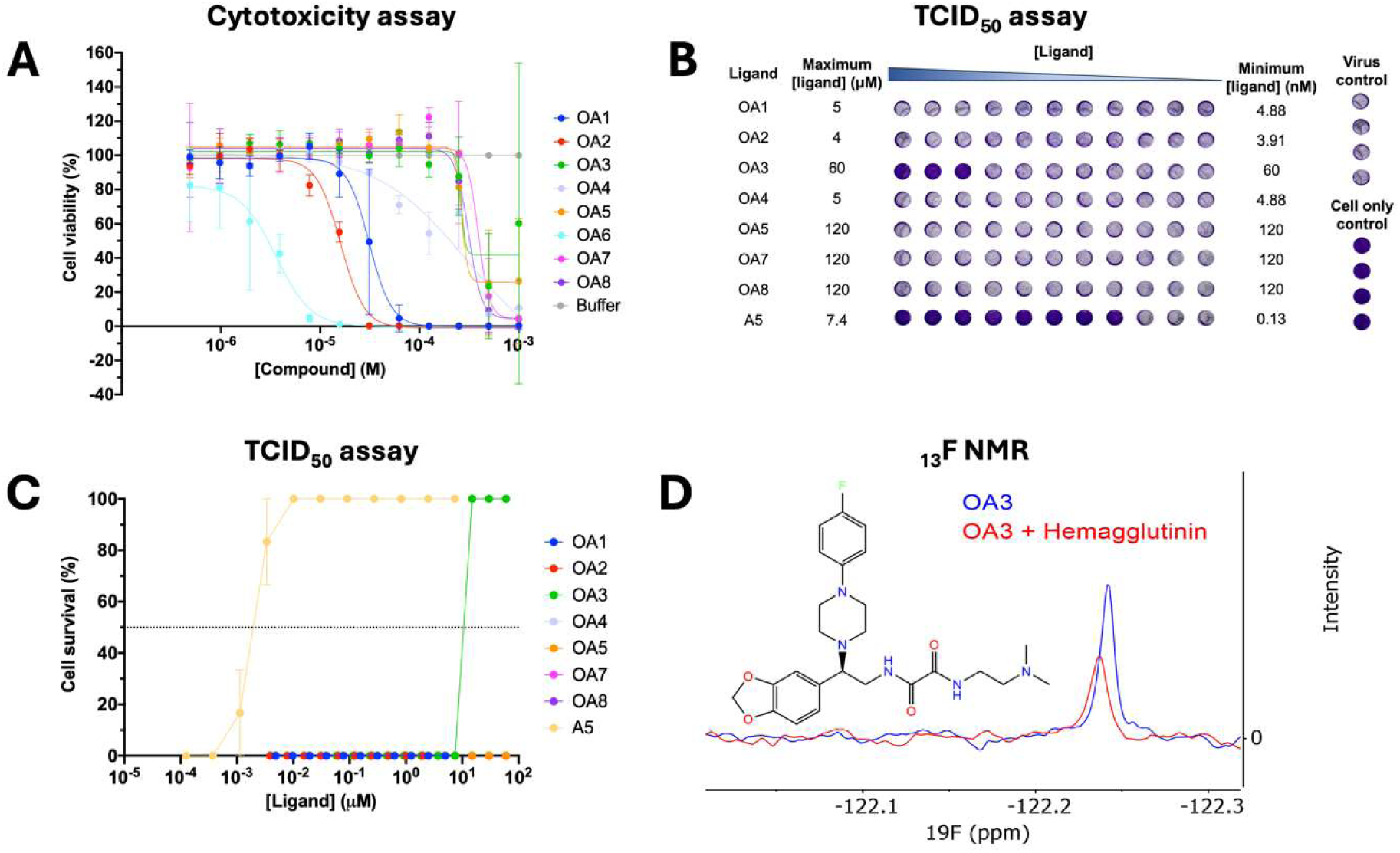
Testing of the OA compound series developed based on an Adhiron which inhibits inffuenza A virus Hemagglutin (HA). A) Cytotoxicity assay in MDCK cells for each compound at a range of concentrations (N=3). B) Representative TCID50 assay plate. C) Combined TCID50 assay results (N=3). D) ^1S^F NMR trace of the Car-Purcell-Meiboom-Gill (CPMG) sequence using a pulse delay of 300 ms for OA3 in the presence (red) and absence (blue) of HA.

The inhibitory properties of the different compounds were assessed against IAV infection via TCID_50_ assay, whereby MDCK cells are infected at a concentration of virus that causes full lysis of the cell monolayer [27]. If a compound inhibits IAV infection at a given concentration, then it will prevent cell lysis, and an intact monolayer would be observed via crystal violet staining. When the OA compounds were screened using the TCID_50_ assay, compound OA3 protected cells against IAV-induced cell death with an IC_50_ value of 11.14 µM (Fig. 4B). Consistent with the initial Adhiron screens, Adhiron A5 protected cells against IAV infection at low nM concentrations [6].

Due to OA3 interfering with the fluorescence wavelengths measured using nanoDSF (data not shown), we employed ligand-based ^19^F NMR CPMG experiments, reasoning that it has limited routes for interference because fluorine atoms are not found naturally in biological material [28]. In CPMG experiments, binding of a fluorinated small molecule to its protein target is indicated by the broadening and reduction of the ^19^F peak height upon addition of the protein target, due to reduced tumbling of the molecule in solution. Therefore, OA3 was screened against trimeric HA. The spectra from the CPMG experiments, when using a 300 ms pulse delay and with HA present, showed a broader and smaller peak, compared to OA3’s native CPMG decay (Fig. 4C). Using 5 CPMG timepoints and data analysis in MNova, a T_2_ value, which is dependent on how quickly OA3 tumbles, can be calculated. In this case, the T_2_ value of OA3 decreased by 69% when HA was present, indicating a reduced tumbling speed attributed to the binding of OA3 to HA. The ^19^F peak is also seen to shift to the left (Fig. 4C), suggesting the fluorine atom is experiencing a different environment when HA is present in the mixture, giving more evidence for binding. Overall, these results illustrate how small molecule modulators of a target protein can be identified based on Adhiron *in silico* models, in the absence of high-resolution structural information.

## Discussion

In this work we demonstrate how Adhirons can be used as starting templates for developing small molecules that mimic their binding regions. In the first example, a high-resolution crystal structure helped to focus virtual screening to the CCHFV NP binding region of NP-Adhiron. The ligand-based virtual screening led to seven small molecules that were tested in both biophysical and biological activity assays, identifying one compound, AF1-02, that demonstrated binding as well as activity against a CCHFV replication assay. By inhibiting the CCHFV MGS assay, AF1-02 was the first example of a biologically active compound derived via mimicking an Adhiron structure. However, it is important to note its cytotoxicity at higher concentration. Therefore, if it was to be taken for hit-to-lead optimisation, removing this cytotoxicity would be key, perhaps by reducing the required dose via improved potency, or by making informed structural changes to reduce cytotoxicity.

In the second example, the lack of high-resolution structure of Adhiron A5 bound to its target HA from IAV meant that ligand-based virtual screening was performed on the entire potential binding region, the Adhiron loops, using computationally generated models. Of the eight compounds purchased, OA3 demonstrated binding to HA and activity against live IAV.

Compared to its template Adhiron, OA3 showed a much lower potency for IAV with an IC_50_ of 11.1 µM vs. 2.0 nM for A5. OA3’s lower potency is not surprising given that OA3 is not an exact mimic of A5 and is much smaller, with A5 having a mass of 11,980 Da and OA3 having a mass of 486 Da. Rather than directly comparing potency via IC_50_ measurements, it is useful to consider ligand efficiency; how effective a ligand is per atom. The binding efficiency index (BEI) is defined as pIC_50_/Molecular mass (kDa), meaning A5 has a BEI of 0.7, whereas OA3 has a higher BEI of 10.2. Therefore, although OA3 is less potent than A5, it is ∼15-fold more efficient and is an initial step towards a lead compound. Even if we only consider the mass of loop one, since the scaffold does not contribute towards Adhiron binding, loop one has a BEI of 7.52, which is still lower than that of OA3.

Although only nanoDSF and NMR results are reported here, both surface plasmon resonance (SPR) spectroscopy and dye-based differential scanning fluorimetry were first tried to measure small molecule binding. However, we found that many small molecules interfered with accurate measurements in these assays, indicating nanoDSF is a useful technique that is both high-throughput and less prone to compound interference. However, nanoDSF was unable to identify binding of OA3 to HA, so ^19^F NMR was employed, indicating a use for a lower throughput technique with even fewer routes for interference. Compounds likely interfered in these assays because as initial hits, they are weak binders which require testing at high concentrations.

Finally, we propose a workflow for the identification of small molecule inhibitors based on previously isolated Adhirons from virtual screening to compound selection and finally *in vitro* evaluation (Fig. 5).

**Figure 5:**
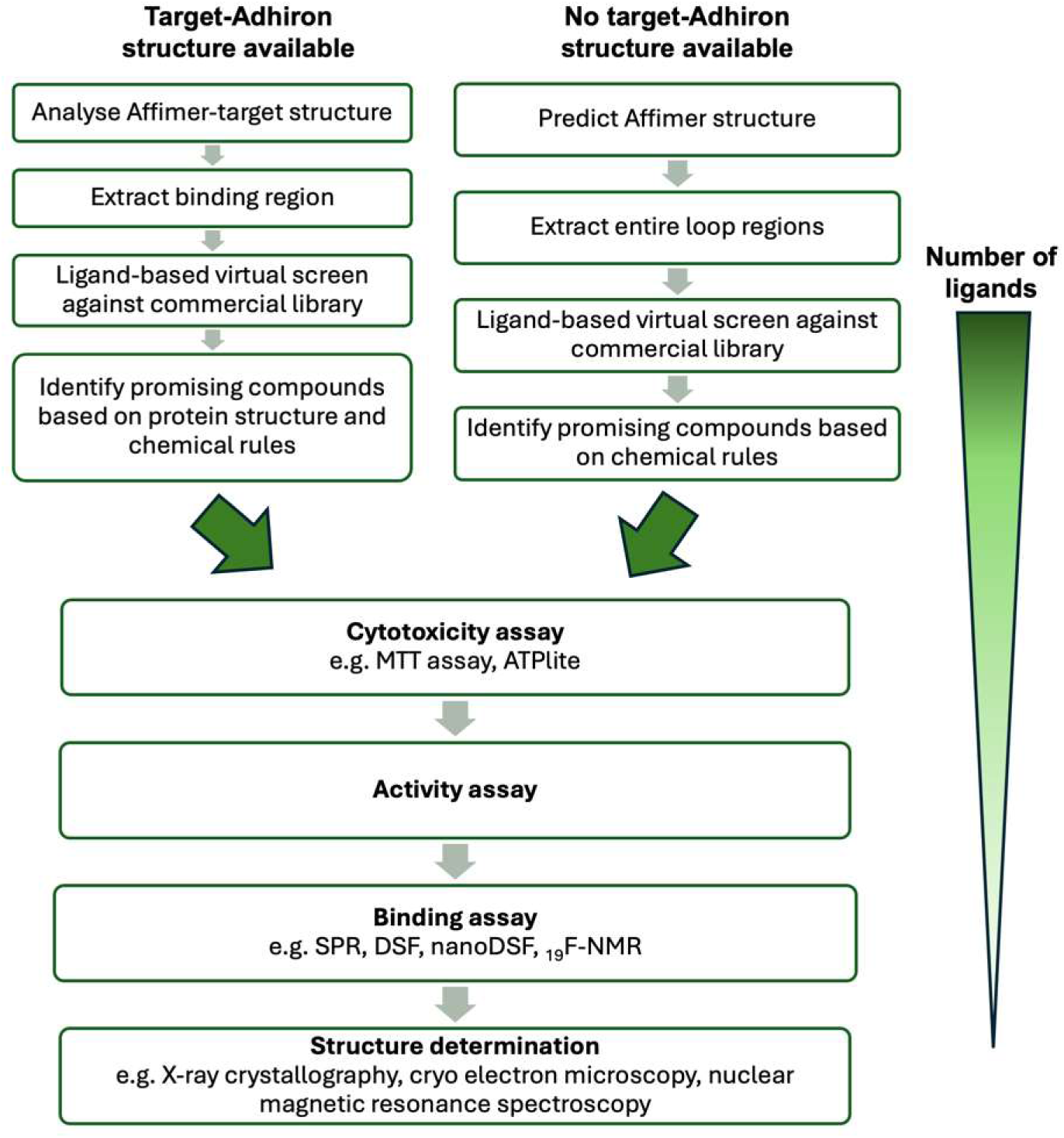
A workflow for generating small molecules mimicking Adhirons.

The virtual screening depends on the availability of a prior Adhiron-bound target structure to focus the screening around the exact Adhiron functional groups to mimic. Although we identified hits both with and without an Adhiron-bound structure to guide the screening, we suspect an experimentally derived structure would improve the hit identification rate and would allow for more rational downstream hit-to-lead development. If no structure is available, homology modelling can be used as was performed here, but moving forward, we recommend AlphaFold 3 for a more accurate prediction and for its ability to predict intermolecular interactions [29].

It is worth noting that the entire loops from some Adhirons like A31 are unsuitable as templates for small molecule development because they contain too many large amino acids, which cannot be easily mimicked by small molecules. In this case, there likely need to be either more efforts to identify the core binding region based on a high-resolution structure or else these Adhirons may need to be disregarded as templates.

For library selection, there are many commercial libraries than can be chosen from. The eMolecules library was selected here as it combines many suppliers into one database. When screened in this work, the eMolecules Screening Compounds database had 12.6 million compounds but now it has 20.3 million compounds [30]. However, there are also many make-on-demand libraries which have a much larger range of compounds, such as the Enamine REAL space database of 64.9 billion possible small molecules and the Enamine xReal space database of 2.4 trillion small molecules that could be synthesised from a stock of building blocks [31].

For the ligand-based virtual screening, we employed ROCS and EON from the OpenEye suite, which worked well, and ROCS allowed parallelisation for use on a high-performance computing cluster. There are even faster shape similarity screening programs which would more easily allow for screening of the billion and trillion sized databased previously mentioned. For example, fastROCS is a GPU-accelerated version of ROCS that allows for screening of one billion compounds in five days on a four GPU cluster or 30 minutes on a cloud-based web server [32].

For compound selection, we recommend the rules we used for the AF1 and OA series. Where there is no Adhiron-target structure, only general rules can be followed to ensure compounds look like the predicted structure of the Adhiron template, such as mimicking the Adhiron backbone shape and overlaying well with at least one side chain. There are also more general rules about compound suitability that can be followed, such as the compound preferably containing a central aromatic group to introduce rigidity and avoiding those with long flexible chains and large hydrophobic groups that may reduce solubility. If guided by a high-resolution Adhiron-bound structure, the compounds should be selected to mimic the important Adhiron functional groups involved in target-binding.

For the screening cascade, we recommend starting with an activity assay to focus on active hits rather than simply binders. Cell-based assays were used here as the target proteins were virus-based, but the assay choice would depend on the target system. Employing cell-based assays means compounds must first be tested for cytotoxicity. Although this is an extra step, cytotoxicity assays are at least high-throughput. Then, active compounds should be tested for binding in high-throughput binding assays such as SPR, DSF or nanoDSF. It should be mentioned that SPR and DSF were employed here, but we observed high compound interference at the concentrations we screened compounds. If these fail, we recommend nanoDSF as a medium-throughput alternative which is less sensitive to interference due to its lack of extrinsic fluorescent dye. ^19^F NMR was used to measure HA-OA3 binding, but it suffers from low throughput. There are of course many other binding assays that could be employed outside of those mentioned here.

An alternative route for this system could be to synthesise bespoke compounds that are better mimics than those found in commercial libraries. While this requires time and chemistry expertise, this method might generate better initial hits, with synthetic routes already in place for structure-activity relationships to be developed.

## Acknowledgements

We thank John N Barr and Darren Tomlinson for useful discussions. Part of this work was undertaken on ARC3, part of the High Performance Computing facilities at the University of Leeds, UK. We thank Dr Andrew Leech at the Department of Biology at the University of York for access to the nanoDSF machine and useful advice. This work was supported by grant PID2023-149259NB-I00, funded by MICIU/AEI/ 10.13039/501100011033 and by “ERDF A way of making Europe” (to J.F.) and Wellcome trust equipment grant 221538/Z/20/Z, which supports the use of the Incucyte live cell imaging platform. A.J.F. thanks the Wellcome Trust for their PhD studentship funding. A.D. thanks The MRC Discovery Medicine North (DiMeN) Doctoral Training Partnership for their PhD studentship funding.

## Methods

### CCHFV NP:NP-Adhiron crystal structure analysis

In order to identify the key Adhiron residues for mimic development, the Adhiron-bound CCHFV NP crystal structure (PDB: 6Z0O) was used as an input for the Protein Interfaces, Surfaces and Assemblies’ service (PISA) hosted at the European Bioinformatics Institute [19]. The interfaces between NP-Adhiron (chain E) loops 1 and 2 and CCHFV NP (chain A) were analysed and the accessible surface area (Å^2^), buried surface area (Å^2^) and solvation energy effect (kcal/mol) was calculated at each residue. The structure was also visually inspected in PyMOL 2.3.2 [33]. Polar interactions between the Adhiron and NP were identified as well as the orientation of binding.

### Ligand-based virtual screening

The 3D structure of DFWKD at NP-Adhiron positions 106-110 within the CCHFV NP:NP-Adhiron crystal structure (PDB: 6Z0O) was extracted into a new model using PyMOL 2.3.2 [33]. The extracted structure was used as a query molecule in the shape similarity search program ROCS 3.2.1.4 [20,21]. The compound library was the eMolecules small molecule library, which had been converted to a maximum of 10 conformers/compound using OMEGA 2.2.0.5 [34,35] giving 12.6 million different conformers in our 3D library. ROCS was run on the high-performance computing cluster ARC3 at the University of Leeds.

The top scoring 10,000 molecules from ROCS were then used as input for the electrostatic similarity search program EON (OpenEye) [36], using the DFWKD structure as the template. The top ∼150 scoring compounds from EON were visually inspected using VIDA (OpenEye). Promising compounds were identified based on the following general criteria:

1) Good mimic of the Adhiron backbone shape.
2) Good mimic of at least one side chain.
3) A central aromatic group to introduce rigidity.
4) Lack of long flexible chains in the molecule.

Additionally, given that the Adhiron-target structure was known and we had detailed information on the interaction, we selected molecules that mimicked at least one hydrogen bond donor or acceptor present in the NP:NP-Adhiron crystal structure. Promising compounds were screened for availability to purchase, then clustered by similarity. A 2D chemical fingerprint was assigned to each compound using Canvas (Schrödinger Release 2025-1: Canvas, Schrödinger, LLC, New York, NY, 2025) [37,38] and they were reduced to the eight compounds with the maximum structural diversity according to the Tanimoto coefficient [39]. Five milligrams of each of the eight compounds were purchased from various suppliers through eMolecules and the compounds were given IDs of AF1-01 – AF1-07.

### Cytotoxicity assay of the AF1 series

BSR-T7 cells were seeded in a 96-well plate and incubated at 37°C and 5% CO_2_ for 16-24 hours until they were ∼80% confluent. Compounds were diluted to 10 µM in 100 µL growth media (DMEM + 1% Pen/Strep + 10% FBS) and applied to cells. The cells were incubated with the compounds for 3 days. Finally, cell viability was measured using the ATPlite kit (Perkin Elmer) according to the manufacturer’s instructions and a FLUOstar OPTIMA fluorescent plate reader (BMG Labtech).

### Mini-genome system assay

BSR-T7 cells were seeded at 9.0 x 10^5^ cells/well in a 6-well plate and the plate was incubated at 37°C and 5% CO2 for 16-24 hours until they were ∼80% confluent. For each well in a 6-well plate, 1.125 µg CCHFV L support plasmid, 375 ng CCHFV N support plasmid and 375 ng S seg UTR-eGFP plasmid were mixed gently in Gibco Opti-MEM reduced serum media (Thermo Fisher Scientific). 2.5 µL room-temperature Trans-IT LT1 reagent (Mirus) per µg plasmid DNA was mixed in with the DNA to give a final volume of 250 µL. For the negative control wells, the L support plasmid was excluded and for the mock transfection, no DNA or Trans-IT was added. The mixture was incubated for 30 minutes at room temperature.

After the incubation, cells were washed in PBS and 1.75 mL fresh growth media (DMEM + 10% FBS + 1% PenStrep) was dispensed onto each well. The transIT-LT1 Reagent:DNA complexes were added drop-wise to the wells and mixed by gentle rocking. The plate was incubated at 37°C in 5% CO_2_ for 24 hours in an Incucyte S3 live-cell imager (Sartorius) to measure GFP expression.

After 24 hours, the cells were trypsinised and re-seeded into wells of a 96-well plate at 3 x 10^4^ cells/well in 50 µL growth media. 50 µL of compound at 20 µM concentration was dispensed onto the cells to give a final compound concentration of 10 µM, with triplicate wells per compound. A DMSO control and media control was included. The plate was incubated at 37°C in 5% CO_2_ for 72 hours in an Incucyte S3 live-cell imager (Sartorius) to measure GFP expression which was normalised to the DMSO control.

### Protein production

Nucleoprotein (NP) from the Baghdad 12 strain of Crimean-Congo hemorrhagic fever virus (CCHFV) and CCHFV NP-Adhiron were produced as described previously [40] [5].

### Nano differential scanning fluorimetry

5 µM NP, and 100 µM compound were diluted to 25 µL with buffer (20 mM Tris pH 7.4, 300 mM NaCl). A 1% DMSO control was used for normalising compound-containing conditions. Tubes were loaded, placed in a Prometheus NT.48 nanoDSF machine. The tubes were heated from 20-95 °C at a rate of 1 °C/minute and the fluorescence was measured using the highest detector energy sensitivity of 100%. The raw fluorescence data from duplicate capillaries were averaged and the derivative of the fluorescence plots was calculated. The inflection points in the derivative plots which corresponded to the target protein with or without ligands was used to identify shifts caused by ligand binding.

### Adhiron A5 homology modelling

Homology models of the HA Adhirons were generated using the I-TASSER server with the default parameters [41]. The template for the modelling was a crystal structure of an Adhiron isolated against the Bcl-xL protein, which was the Adhiron for which a structure was available with the highest sequence similarity to the HA Adhirons [24] (PDB: 6HJL). The loop regions were extracted from the homology models using PyMOL 2.3.2 [33] and used as inputs for ROCS [20,21].

### Ligand-based virtual screening

Small molecule mimics of the A5 loops were identified using shape similarity program ROCS using the same workflow outlined for NP-Adhiron, except only the top 1000 conformers from the ROCS run were used as input for EON [20]. The general rules for identifying promising compounds were the same as those for the AF1 mimics of NP-Adhiron. However, due to the lack of a high-resolution HA-A5 structure we also aimed to identify small molecules that contained at least one potential hydrogen bond donor or acceptor (since the exact groups involved in hydrogen bonding were unknown). The compounds were selected for purchase according to availability and to represent as many pharmacophores as possible and were given the IDs OA1-OA8.

### Cytotoxicity assay of OA series

MDCK cells were seeded at 0.5 x 10^5^ cells/well in a 96-well plate and incubated at 37°C and 5% CO_2_ for 16-24 hours until they were ∼80% confluent. To test for cytotoxicity of a range of concentrations, the OA compounds were serially diluted 1 in 2 in infection media (DMEM, 0.1% TPCK trypsin) across a 96-well plate from 1 mM to 488 nM and dispensed on to the MDCK cells. The cells were incubated with the compounds for 3 days, after which the cell viability was measured using the ATPlite kit (Perkin Elmer) according to the manufacturer’s instructions and a FLUOstar OPTIMA fluorescent plate reader (BMG Labtech).

### TCID_50_ assay

TCID_50_ assay followed the method used to originally screen A5 [6]. MDCK cells were seeded at 5 x 10^4^ cells/well in a 96-well plate and incubated at 37 °C in 5% CO_2_ for 16-24 hours until they were ∼90% confluent. Starting at 2X the highest compound concentration which led to a >95% cell viability in the cytotoxicity assay, the OA compounds were serially diluted 1 in 2 in infection media (DMEM, 0.1% TPCK trypsin) across a 96-well plate, leaving 50 μL 2X final concentrations. 50 μL influenza virus (A/Aichi/1968/(H3N2)) at 200x the TCID_50_ value for the virus stock was added to each well and mixed 1:1 with the drug dilutions.

Compound and virus were incubated for 45 minutes then applied to the MDCK cells which had been washed with PBS. Cell only wells containing no compound or virus were included as a control along with virus only wells, which contained virus but no compound. After 72 hours, the drug/virus mixture was aspirated off the cells and the cells were washed with PBS, fixed with 4% paraformaldehyde then stained with crystal violet solution for 15 minutes. Finally, the cells were washed with water and imaged.

### NMR experiments

NMR experiments were performed using a 500 MHz four-channel Bruker AV-NEO NMR spectrometer equipped with 5mm TBO probe. All samples were prepared in Eppendorf tubes and transferred to a 5 mm Wilmad 528-PP NMR tube. 2.25 µL of OA3 and 2.25 µL of a control compound ((2,6-Difluorophenyl)-4-thiomorpholinylmethanone) (both at 10 mM stock in DMSO-d^6^) were added to 396 µL of PBS (136.80 mM NaCl, 10 mM Na_2_HPO_4_, 1.98 mM KH_2_PO_4_, 2.68 mM KCl). 49.5 µL of trimeric HA (stock solution of 1 g/L) was added along with 50 µL of D_2_O (10 % of the final volume). In control runs absent of HA, 49.5 µL of PBS was added instead of trimeric HA. This gave final concentrations of each compound as 45 µM and trimeric HA at 0.5 µM giving a ratio of 90:1 of ligand to trimeric protein. Car-Purcell-Meiboom-Gill (CPMG) experiments used 2048 scans for each sample at 5 different relaxation delays (10, 50, 100, 200, 300 ms). Data was processed using MNova V15 to calculate T_2_ values, in addition to generate figures.

## Author Contributions

**Conceptualization -** Alex J. Flynn, Stephen P. Muench, Juan Fontana, Katie Simmons

**Investigation -** Alex J. Flynn, Alex Derry, Oliver Debski-Antoniak, Katie Simmons

**Funding Acquisition** Alex J. Flynn, Stephen P. Muench, Juan Fontana, Katie Simmons

**Methodology -** Alex J. Flynn, Stephen P. Muench, Juan Fontana, Katie Simmons

**Supervision -** Stephen P. Muench, Juan Fontana, Katie Simmons

**Writing – Original Draft Preparation -** Alex J. Flynn, Stephen P. Muench, Juan Fontana, Katie Simmons

**Writing – Review G Editing -** Alex J. Flynn, Alex Derry, Oliver Debski-Antoniak, Stephen P. Muench, Juan Fontana, Katie Simmons

## Supplemental material

**Supplementary Figure 1:**
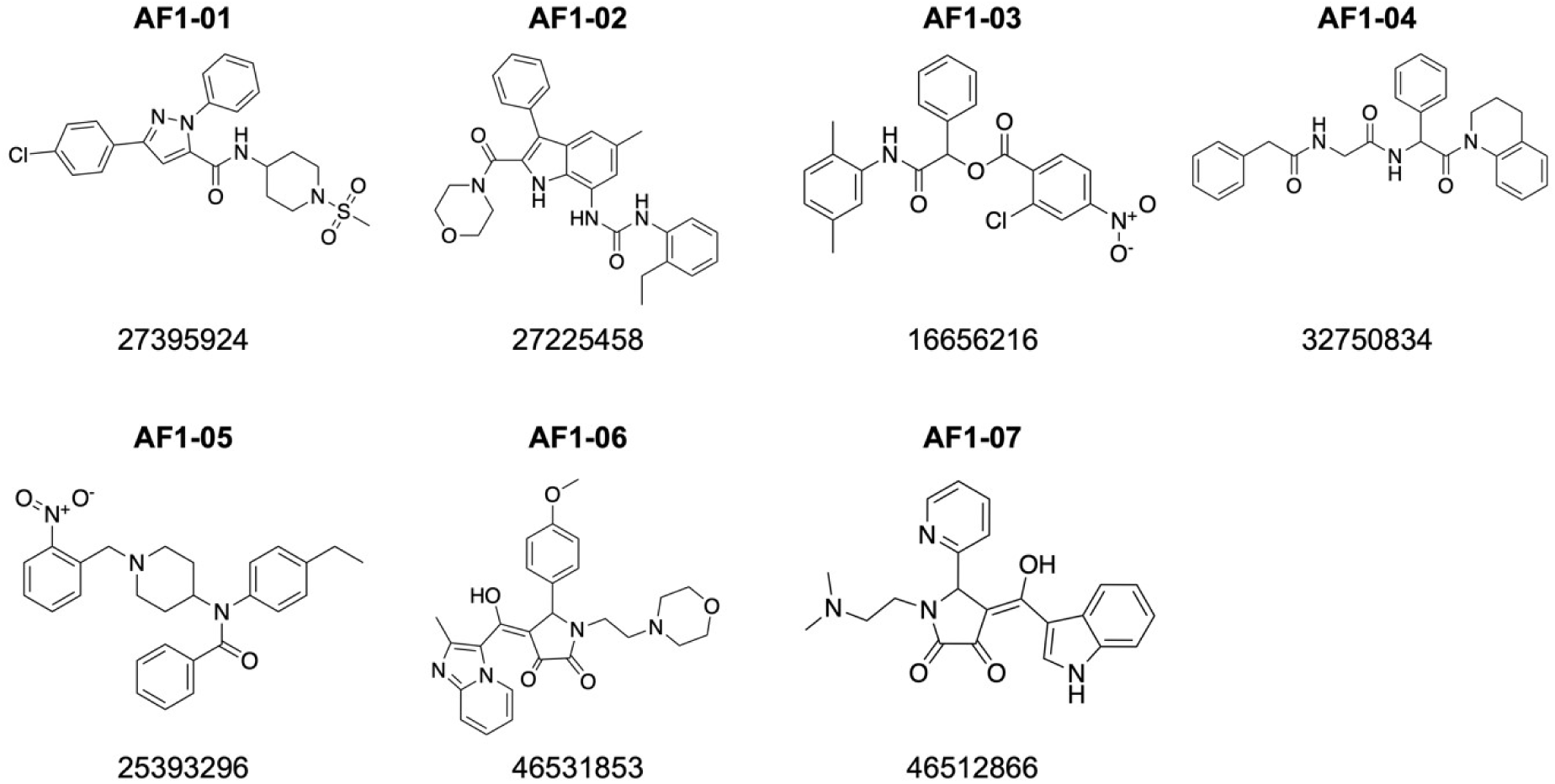
The AF1 compounds were generated as mimics of NP-Adhiron, an Adhiron inhibitor of CCHFV NP. For each compound the AF1 ID (top) and its eMolecules ID number (bottom) are shown.

**Supplementary Figure 2:**
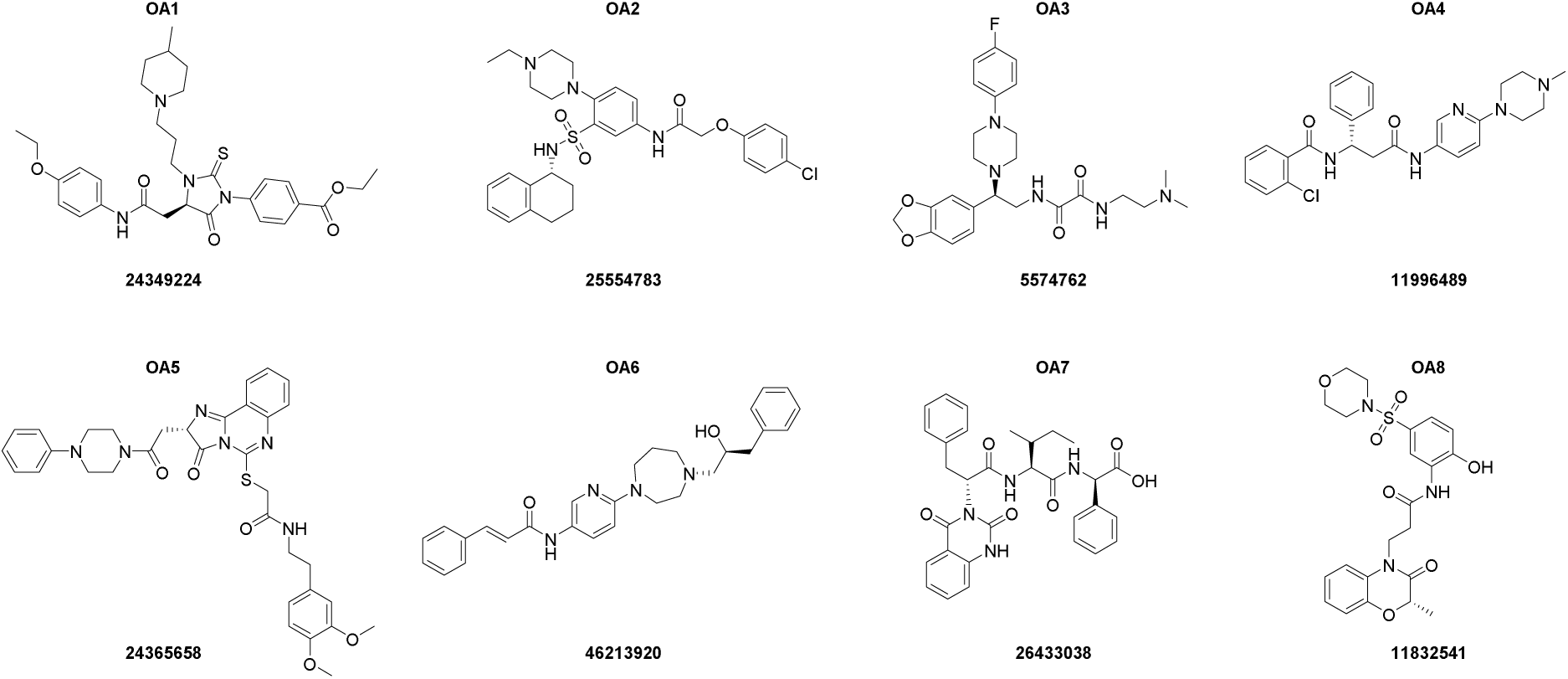
The OA compounds were generated as mimics of A5, an Adhiron inhibitor of IAV HA. For each compound the OA ID (top) and its eMolecules compound library ID number (bottom) are shown.

